# Placental and fetal characteristics of the Ohia mouse line recapitulate outcomes in human hypoplastic left heart syndrome

**DOI:** 10.1101/2020.08.17.252387

**Authors:** Rebecca L Wilson, Weston Troja, Jennifer Courtney, Alyssa Williams, Helen N Jones

## Abstract

Congenital heart defects (CHDs) are the most common birth defect worldwide. The morbidity and mortality associated with these defects is compounded by increased frequency of fetal growth abnormalities. In the Ohia mouse model of hypoplastic left heart syndrome (HLHS), the double homozygous genotype is embryonically lethal at mid-pregnancy; a time in which optimal establishment of the placenta is crucial to fetal survival. We aimed to characterize placental and fetal growth and development in the double heterozygous genotype (*Sap130*^*m/+*^*Pcdha9*^*m/+*^) to determine whether the genetic mutations associated with HLHS in the Ohia mouse also affect the placenta. The frequency of fetuses with reduced weight near term was shifted in the *Sap130*^*m/+*^*Pcdha9*^*m/+*^ fetuses compared to wildtype. This shift in fetal weight distribution in the *Sap130*^*m/+*^*Pcdha9*^*m/+*^ fetuses was associated with reduced labyrinth region area (P<0.001) and reduced fetal capillary density (P<0.001) in the placentas. Positive correlations were observed between fetal weight and placenta mRNA expression of several nutrient transporters in the *Sap130*^*m/+*^*Pcdha9*^*m/+*^ fetuses but not observed in the wildtype. Increased protein expression of Slc7a5 (P<0.05) and Slc7a8 (P<0.05) was also found in the placenta of *Sap130*^*m/+*^*Pcdha9*^*m/+*^ fetuses. This data shows, despite a potential compensatory mechanism to increase nutrient transport, abhorrent placental vascularization leads to inadequate fetal growth in the Ohia mouse model. Such differences are similar to findings in studies of human placentas and highlights the importance of this mouse model in continuing to understand the developmental links and disruptions to the heart-placenta axis.

## Background

Congenital Heart Defects (CHDs) are the most common birth defect in the Unites States. The high risk of morbidity and mortality associated with CHD is compounded by increased frequency of low birth weight (LBW; <2500 grams), small for gestational age (SGA; <10th percentile), and microcephaly (<3rd percentile) in these newborns [1]. One such defect is Hypoplastic Left Heart Syndrome (HLHS) and is characterized by the inability of the left ventricle to provide adequate systemic circulation [2] and is fatal shortly after birth without surgical intervention. The causes of HLHS remain unknown, but considerable evidence has demonstrated both genetic and environmental factors [3, 4]. Newborns with HLHS are critically ill and require multiple reconstructive surgeries early in life. In addition to brain abnormalities, which occur in approximately half of newborns with HLHS, growth abnormalities at birth are common [5, 6]. LBW and SGA are major risk factors for stage I Norwood palliative surgery, death and major complications following stage I palliative surgery [7, 8].

Currently, there is no clear link between fetal growth abnormalities and HLHS; however, fetal genetic factors, maternal health conditions and placental abnormalities have been implicated as contributing factors. The placenta has numerous functions that are crucial to supporting normal fetal growth including nutrient transfer to the developing fetus [9]. Thus, abnormal placentation, development and function can negatively impact fetal growth leading to fetal growth restriction (FGR). The paucity of knowledge regarding the causes of FGR in pregnancies with fetuses that develop HLHS is being mitigated by the generation of the Ohia mouse model of HLHS [10]. The HLHS-like phenotype in the Ohia mice arises from mutations in 2 genes: *Sap130* and *Pcdha9* with homozygotes being either embryonic lethal before or at mid-gestation, or rarely survive to late gestation and demonstrate severe cardiac defects such as hypoplasia of the left ventricle, aorta and mitral valve [11].

The timing at which embryonic lethality occurs in the Ohia mouse model suggests the mutation has a significant impact on placental development. Placental development in the mouse can be divided into 2 broad stages [12, 13]. The first stage of placental development last from implantation to mid-pregnancy (gestational day 9.5) and is characterized by trophoblast invasion and differentiation to form the mature mouse placenta. Fetal growth is supported by the yolk sac while the labyrinth develops to coordinate placental nutrient transport. The second stage is from mid-pregnancy onwards, where maternal blood flows into the mature labyrinth to support fetal growth. It is widely accepted that the switch at mid-pregnancy creates a developmental bottleneck leading to embryonic death in numerous mouse mutants [12, 14]. In the Ohia mouse, extensive investigations have been conducted into heart development however, other aspects of fetal and placental development have not been investigated. Therefore, we aimed to determine whether the genetic mutations associated with HLHS in this mouse model also disrupt placental development and function.

## Methods

### Mouse procedures

C57Bl/6J-b2b635Clo/J (Ohia) mice were acquired from Jackson Laboratory (*Bar Harbor, ME*). All animal procedures were performed under protocol 2015-0087 and approved by the Institutional Animal Care and Use Committee of Cincinnati Children’s Hospital Medical Center. Animals were housed in individual cages (1-4 per cage), maintained in a 12-hour light/dark cycle, and food and water provided *ad libitum*. Ohia males double heterozygous for *Sap130* and *Pcdha9* (*Sap130*^*m/+*^*Pcdha9*^*m/+*^) were time mated with *Sap130*^*m/+*^ *Pcdha9*^*m/+*^ Ohia females for 12-hours overnight. At gestational day (GD) 18.5, females (n=4) were euthanized by CO_2_ inhalation followed by cervical dislocation, and placentas and fetuses were weighed. Maternal organs, placentas, and fetal tissue were then either snap frozen or fixed in 10% formalin for genotyping and further analysis.

### Genotyping

Genomic DNA (gDNA) was extracted from fetal tails and adult ear clippings according to the HotSHOT protocol [15]. Oligonucleotide primers were designed using Primer3 to amplify ∼730bp regions surrounding *Sap130* and *Pcdha9* single nucleotide polymorphisms (SNPs) as identified by Liu et al. [11]. Primer sequences were as follows: Sap130 (PCR) F: CCAGTGTGAAGTGAGTCAGCA, R: AGCACCATGTGCCTGTAATCT; Pcdha9 (PCR) F: CCTATGTTGATCGCCACTGC, R: CACCCTGAACACAAGTTCTATTGG; Sap130 (Sequencing) F: TTTTTACCCTCTTTTCAGGATCT, R: TCTGGGACACACATTTGTCAT; Pcdha9 (Sequencing) F: ACCTGATCATTGCCATTTGC, R: AAGTCAAATTCAGAGTTGCTT. gDNA regions were amplified via PCR using 1/150th gDNA and 300nM forward and reverse primers in a 50 μL FastStart PCR Master Mix reaction (Roche). PCR products were purified using QIAquick PCR Purification Kit (*Qiagen*). Purity and concentration was determined using agarose gel electrophoresis. To assess SNPs, 30 ng of purified PCR product and 10 pmols of forward and reverse sequencing primers in a 12 μl reaction were submitted to the DNA Core at Cincinnati Children’s Hospital Medical Center for Sanger sequencing. SNPs for *Sap130* and *Pcdha9* were confirmed using FinchTV (*Geospiza Inc*).

### Placental Histology and Immunohistochemistry

Placental tissues were fixed in 10% formalin for 24 hours at 4°C immediately after dissection. Tissues were embedded in paraffin, cut into 5μm serial sections, deparaffinized in xylene, and rehydrated. To assess physical structure of Ohia placentas, sections were stained with Hematoxylin and Eosin (H&E) following standard protocol and double label immunohistochemistry (IHC) was performed in order to assess the volume densities of trophoblast and fetal capillary cells in the labyrinth following the protocol outlined in [16]. IHC analysis of the protein expression of nutrient transporters was performed as follows: sections were incubated in 1X sodium citrate (pH 6.0) target retrieval solution (*Dako*) at 95°C for 30 minutes. Endogenous peroxidase activity was blocked by incubation in 3% hydrogen peroxide, followed by incubation with a 10% goat serum and 1% BSA protein block to limit non-specific antibody binding. Nutrient transporters Slc2A1 (1:750; ab15309, *Abcam*), Slc7a5 (1:100; ab85226, *Abcam*) and Slc7a8 (1:250; ab75610, *Abcam*) were diluted and applied to sections overnight at 4°C. Antibody binding was then amplified by incubating with goat anti-rabbit secondary antibody (1:200, *Vector Laboratories*), followed by incubation with the ABC reagent (*Vector Laboratories*), and visualized using DAB (3,3′-Diaminobenzidine; *Dako*). Sections were counterstained with hematoxylin, dehydrated, and mounted. Histological examination was conducted by light microscopy using the Axioscan (*Zeiss*). DAB staining intensity of nutrient transporters was analyzed using ImageJ as described [17] on 5 randomly selected images of the placental labyrinth at 10x magnification, per placenta section.

### Placental Gene Expression

Total RNA was isolated from snap-frozen whole placentas using the FastPrep-24 Classic bead beater homogenizer (*MP Biomedicals*) and the RNAeasy Mini Kit (*Qiagen*) per manufacturer’s protocol. 1 μg of total RNA was reverse transcribed into cDNA according to the High Capacity cDNA Reverse Transcription kit protocol (*Applied Biosystems*). For angiogenic factor and nutrient transport analysis, oligonucleotide primers for SYBR Green assays were designed to span intron/exon boundaries, aligned against murine genome via Primer-BLAST to ensure specificity. Primer sequences are as followed: *Angiopoietin 1* (*Ang1*) F: GCACGAAGGATGCTGATAAC, R: AACCACCAACCTCCTGTTAG; *Ang2* F: GCACAAAGGATTCGGACAAT, R: AAGGACCACATGCGTCAAA; *Vascular endothelial growth factor α* (*Vegfα*) F: TTAAACGAACGTACTTGCAGATG, R: AGAGGTCTGGTTCCCGAA; *Placenta growth factor* (*Pgf*) F: GACCTATTCTGGAGACGACA, R: GGTTCCTCAGTCTGTGAGTT; *Kinase Insert Domain Receptor* (*Kdr*) F: CGTTGTACAAATGTGAAGC, R: CACAGTAATTTCAGGACCC; *Fms Related Receptor Tyrosine Kinase 1* (*Flt1*) F: TGACGGTCATAGAAGGAACA, R: TAGTTGGGATAGGGAGCCA; *Slc2A1* F: TGTGCTCATGACCATCGC, R: AAGGCCACAAAGCCAAAGAT; *Slc2A3* F: CATTGTCCTCCAGCTGTCTC, R: GCGTCCTTGAAGATTCCTGT; Slc38A1 F: GGACGGAGATAAAGGCACTC, R: CAGAGGGATGCTGATCAAGG; *Slc38A2* F: TCCTTGGGCTTTCTTATGCC, R: TTGACACGAACGTCAAGAGA; *Slc7A5* F: ATTCAAGAAGCCTGAGCTGG, R: GCAGGCCAGGATAAAGAACA; Slc7A8 F: AAGAAAGAGATCGGATTGGT, R: TTTCGGTGAGACAAAGATTC; *Rps20* F: GCTGGAGAAGGTTTGTGCG, R: AGTGATTCTCAAAGTCTTGGTAGGC. PCR reactions were then performed using 300 nM reverse and forward primers, 1/40^th^ of the cDNA template in a 25 μL reaction of SYBR Green PCR Master Mix. Gene expression assays were completed in duplicate and normalized to murine *Rps20* using Applied Biosystems StepOne-Plus Real-Time PCR System. Relative expression was calculated using the Comparative Ct (ΔΔCt) method.

### Statistics and data presentation

Data was analyzed using SPPS statistics 26 and graphed using Prism (v8.0.1). Generalized linear modelling was used to calculate p-values with genotype considered a main effect and litter size a covariate. Fetal and placental weight distributions were calculated using a Gaussian non-linear model. Correlations between placenta nutrient transporter mRNA expression and fetal weight were calculated using Spearman’s method. All data are represented as estimate marginal mean + standard error unless otherwise stated and deemed significant with P<0.05.

## Results

### Fetal weight distribution is left-shifted in the Ohia heterozygote and associated with abhorrent placenta morphology

No double homozygote mice were recovered at GD18.5 and therefore, no placentas could be assessed. At GD19.5, mean fetal weight remained at wildtype levels in the double heterozygote genotype (Figure 1A) however, fetal weight distribution was shifted towards lower fetal weights (Figure 1B). Within the double heterozygous genotype, mean placental weight as well as placental weight distribution was similar between the wildtype and double heterozygotes (Figure 1C and 1D). The increased frequency of lower fetal weight double heterozygotes was associated with changes to the placenta microstructure. Placental morphology was highly variable between double heterozygote genotypes but demonstrated abnormal architecture in comparison to wildtype littermates (Figure 2A). Whilst there was no difference in total mid-sagittal cross-sectional area between double heterozygous and wildtype placentas (Figure 2B), there was an increase in junctional area in the double heterozygous placentas (Figure 2C) and as a result, a reduction in labyrinth area when compared to wildtype placentas (Figure 2D). Additionally, there was a reduction in fetal capillary density in the double heterozygote placentas (Figure 3A and 3B).

**Figure 1.**
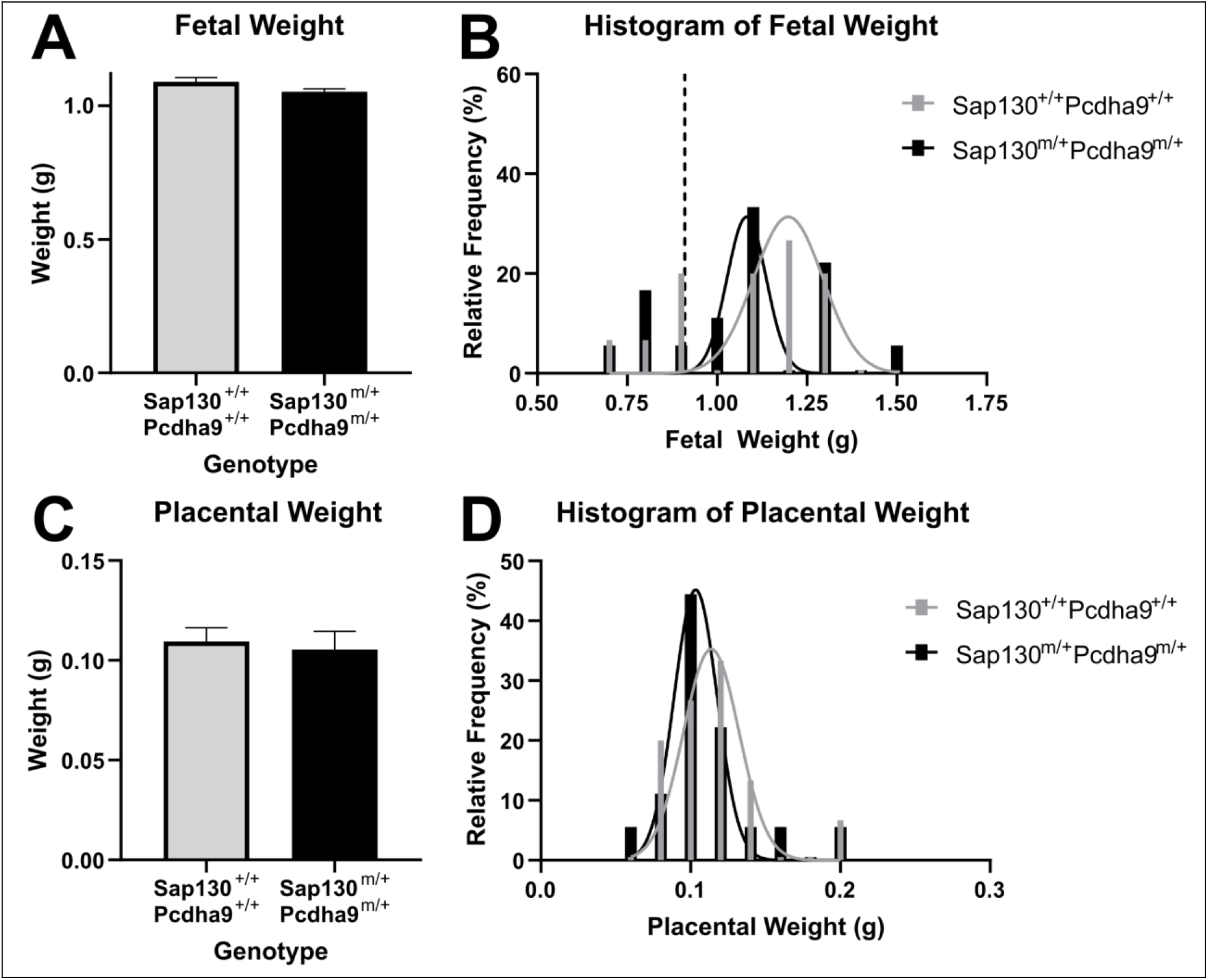
Fetal and placenta weight in Ohia mouse at GD18.5. Mean fetal weight was not different between double heterozygous (*Sap130*^*m/+*^ *Pcdha9*^*m/+*^) and wildtype (*Sap130*^*+/+*^ *Pcdha9*^*+/+*^)(**A**) however, there was a shift in fetal weight distribution toward lower fetal weight in the *Sap130*^*m/+*^ *Pcdha9*^*m/+*^ genotype (**B**). Neither mean placenta weight (**C**) nor placenta weight distribution (**D**) was different between the double heterozygous and wildtype genotypes. Data are estimated marginal mean + standard error, n=4 females (15 wildtype and 18 double heterozygous fetuses), analyzed using generalized linear modelling and including litter size as a covariate. Weight distributions were calculated using Gaussian nonlinear regression analysis.

**Figure 2.**
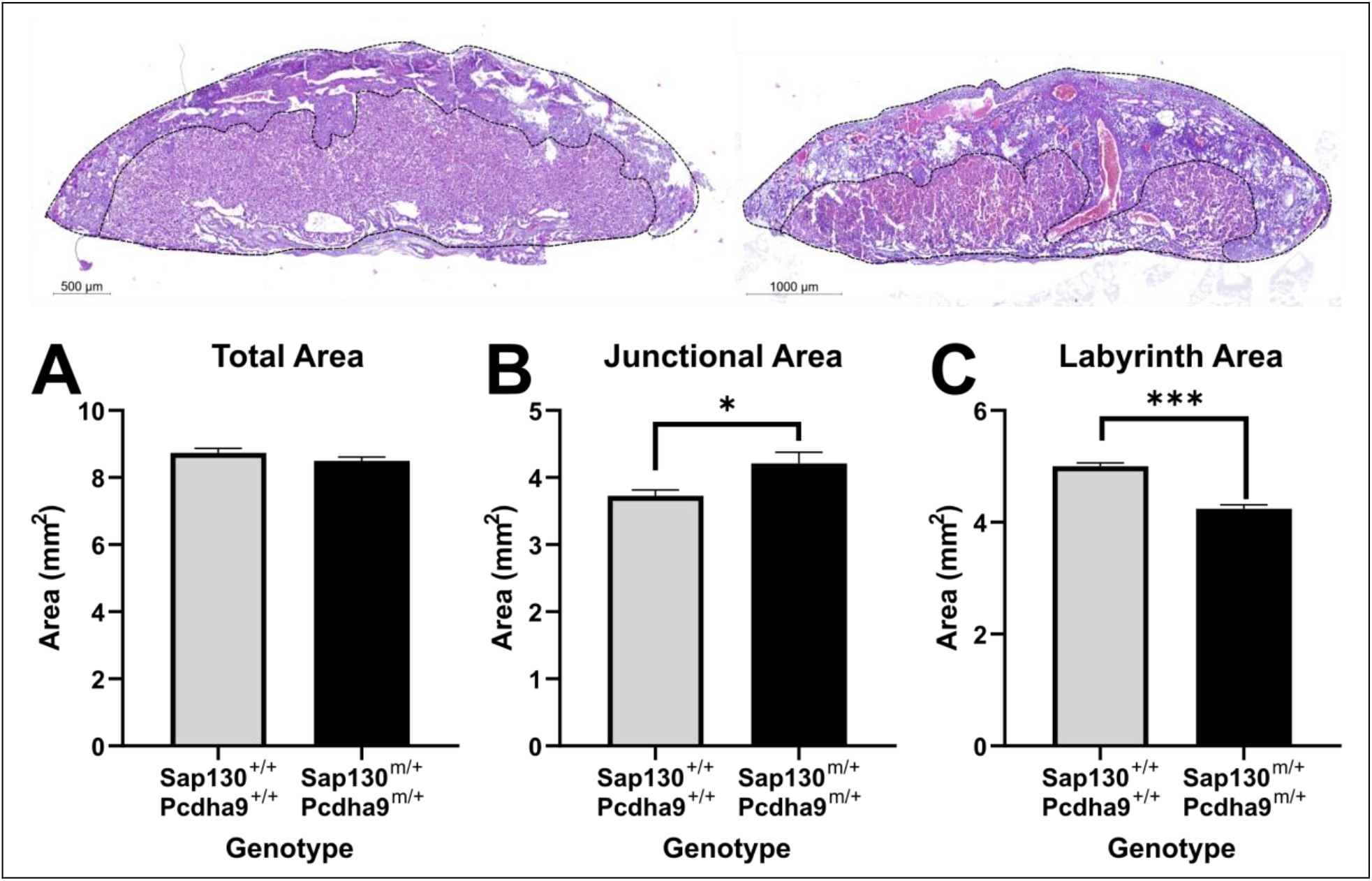
Gross placental structure of the Ohia mouse at GD18.5. Compared to the wildtype (*Sap130*^*+/+*^ *Pcdha9*^*+/+*^; Left representative image) genotype, placental structure in the double heterozygous genotype (*Sap130*^*m/+*^ *Pcdha9*^*m/+*^; Right representative image) was highly variable and exhibited abnormalities within the junctional and labyrinth regions. Whilst there was no difference in the total mid-sagittal area (**A**), there was an increase in junctional region area in the double heterozygous placentas when compared to the wildtype placentas (**B**). As a result, there was a decrease in labyrinth region area in the double heterozygous placentas compared to wildtype (**C**). Data are estimates marginal mean + standard error, n=3 females (9 wildtype and 14 double heterozygous), analyzed using generalized linear modelling and including litter size as a covariate. *P<0.05; ***P<0.001

**Figure 3.**
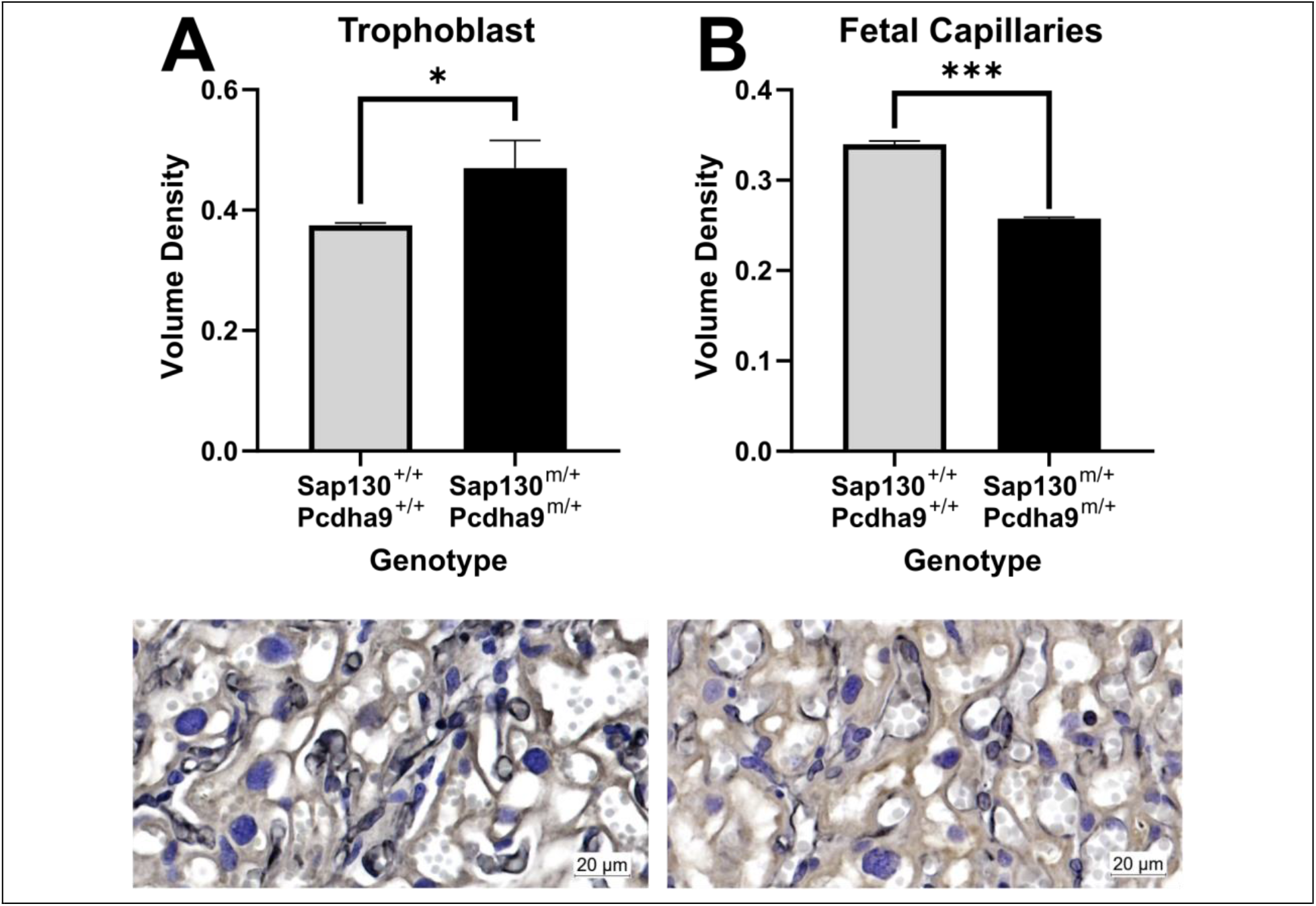
Placental microstructure of the Ohia mouse at GD18.5. Compared to the wildtype (*Sap130*^*+/+*^ *Pcdha9*^*+/+*^) genotype, there was an increased volume density of trophoblast cells in the labyrinth of the double heterozygous genotype (*Sap130*^*m/+*^ *Pcdha9*^*m/+*^) (**A**). As a result, the volume density of fetal capillaries was reduced in the double heterozygous genotype (**B**). Representative image of labyrinth double label immunohistochemistry identifying trophoblast (brown) and fetal capillaries (black) in wildtype (Left) and double heterozygous (Right) placentas. Data are estimates marginal mean + standard error, n=3 females (9 wildtype and 14 double heterozygous), analyzed using generalized linear modelling and including litter size as a covariate. *P<0.05; ***P<0.001

### Reduced placental capillary volume density was associated with reduced placenta mRNA expression of angiogenic factors Pgf and sFlt1

Changes in placenta microstructure of the Ohia mouse suggested reduced fetal angiogenesis. Compared to wildtype genotype, placenta mRNA expression of *Pgf, Flt1* and *Kdr* was reduced in the double heterozygotes (Figure 4A-C). There was no difference in the expression of *Vegfα, Ang1* and *Ang2* between the different genotypes (Figure 4D-F).

**Figure 4.**
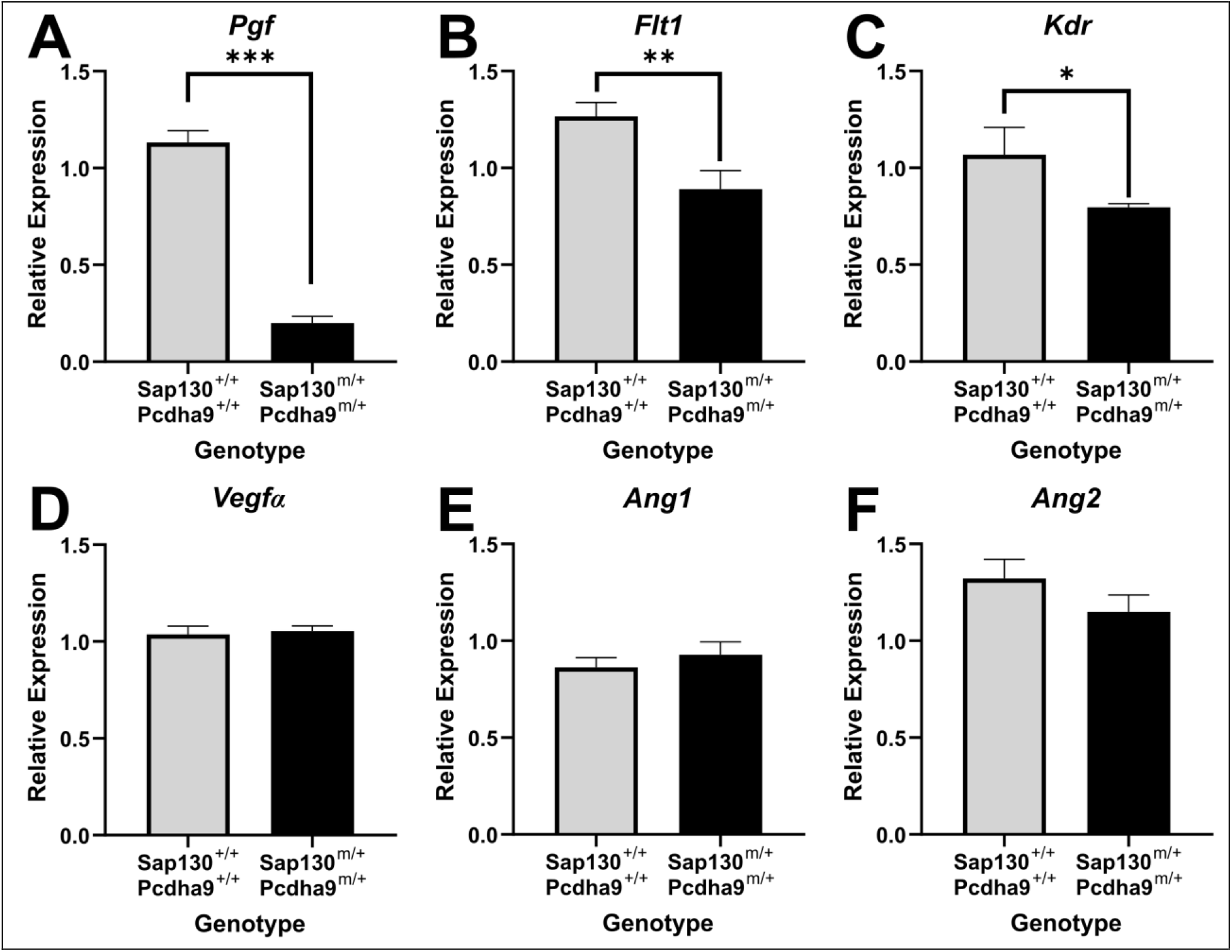
mRNA expression of angiogenic factors in the placenta of the Ohia mouse. Compared to wildtype (*Sap130*^*+/+*^ *Pcdha9*^*+/+*^), expression of placenta growth factor (*Pgf*; **A**), fms-like tyrosine kinase-1 (*Flt1*; **B**) and kinase insert domain receptor (*Kdr*; **C**) was reduced in the double heterozygous (*Sap130*^*-/+*^ *Pcdha9*^*-/+*^) placenta. There was no difference in the expression of vascular endothelial growth factor α (*Vegfα*; **D**), angiopoietin 1 (*Ang 1*; **E**) or 2 (*Ang2*; **F**). Data are estimated marginal mean + standard error, n=3 females (6 wildtype and 7 double heterozygous placentas), analyzed using generalized linear modelling and including litter size as a covariate. *P<0.05; **P<0.01; ***P<0.001

### Placenta nutrient transporter expression is increased in the Ohia heterozygote indicative of a compensatory response

Normally, the anatomy of the placenta is highly vascular with complex villous structures that optimize nutrient, oxygen and waste exchange. Whilst mean mRNA expression of nutrient transporters in the placentas of wildtype and double heterozygous genotypes was not different (Supplemental Figure 1), correlations between fetal weight and placenta mRNA expression of several nutrient transporters demonstrated positive correlations in the double heterozygous genotypes but not in the wildtype (Table 1). More specifically, there was a positive correlation between fetal weight and mRNA expression of *Slc2A1, Slc7A5* and *Slc7a8* in the double heterozygous genotype but a negative correlation with fetal weight in the wildtype genotype. Analysis of protein expression showed increased staining intensity of Slc7a5 (Figure 5A) and Slc7a8 (Figure 5B) in the double heterozygous placentas when compared to the wildtype whilst there was no difference in the expression of Slc2a1 (Figure 5C).

**Table 1.** Correlation between fetal weight and mRNA expression of nutrient transporters in the placenta of wildtype (*Sap130*^*+/+*^ *Pcdha9*^*+/+*^) and double heterozygous (*Sap130*^*m/+*^ *Pcdha9*^*m/+*^) Ohia mice

|  | *Sap130<sup>+/+</sup> Pcdha9<sup>+/+</sup>* | *Sap130<sup>m/+</sup> Pcdha9<sup>m/+</sup>* |
| --- | --- | --- |
|  | Spearman r value | Spearman r value |
| *Slc2A1* | -0.0857 | 0.643 |
| *Slc2A3* | 0.429 | 0.679 |
| *Slc38A1* | 0.543 | 0.321 |
| *Slc38A2* | 0.0857 | 0.643 |
| *Slc7A5* | -0.200 | 0.607 |
| *Slc7A8* | -0.200 | 0.789 |

**Figure 5.**
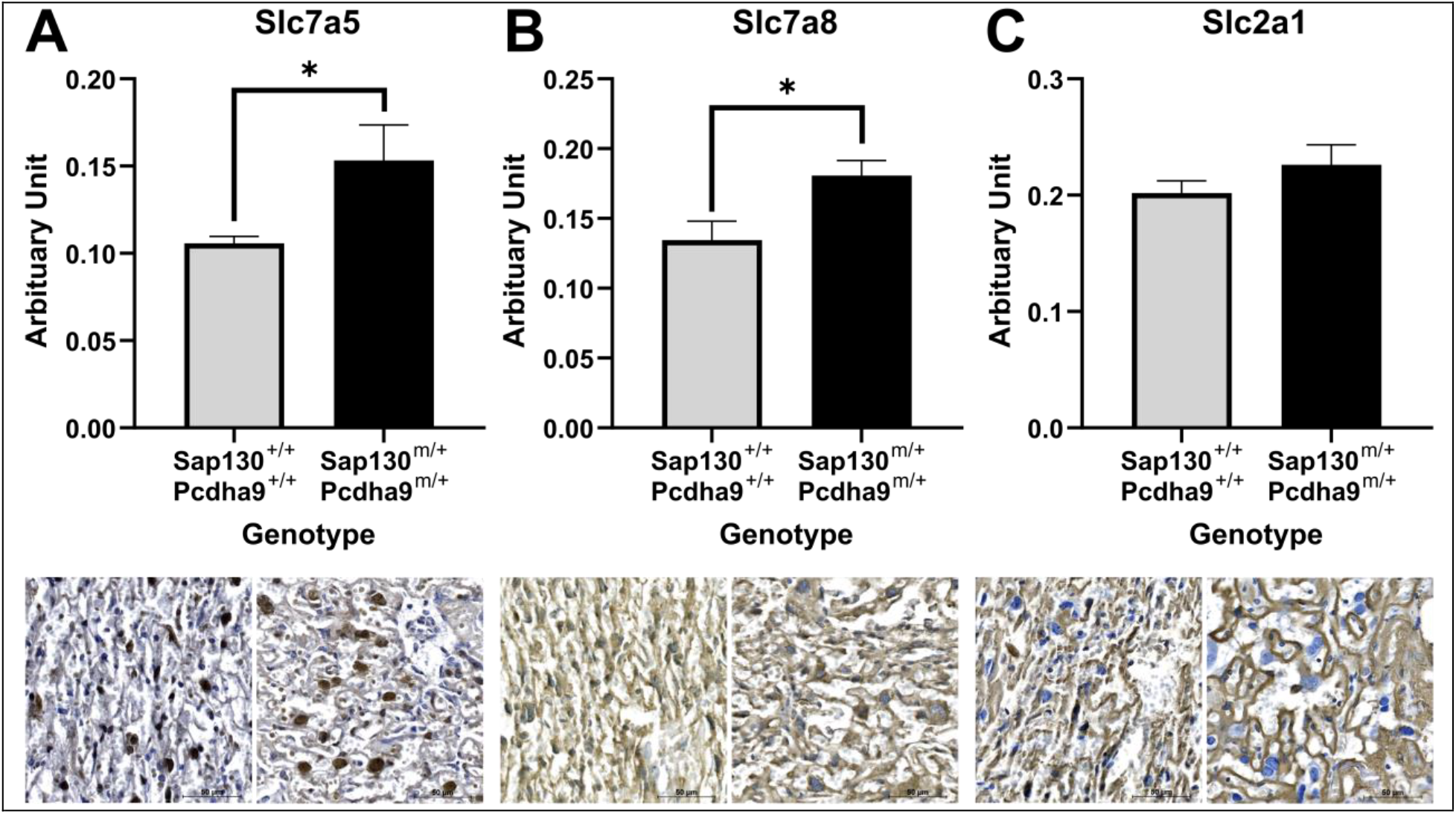
Nutrient transporter expression in the placenta of the Ohia mouse at GD18.5. Compared to the wildtype (*Sap130*^*+/+*^ *Pcdha9*^*+/+*^) genotype, there was an increased staining intensity of nutrient transporter Slc7a5 (A) and Slc7a8 (B) in the labyrinth of the double heterozygous genotype (*Sap130*^*m/+*^ *Pcdha9*^*m/+*^). There was no difference in staining intensity of Slc2a1 between double heterozygous and wildtype placentas (C). Representative images of nutrient transporters in wildtype (Left) and double heterozygous (Right) placentas. Data are estimates marginal mean + standard error, n=3 females (9 wildtype and 14 double heterozygous), analyzed using generalized linear modelling and including litter size as a covariate. *P<0.05

## Discussion

In humans, Congenital Heart Defects such as HLHS are often associated with poor fetal growth that is likely contributed to by inappropriate placental development, growth and function. Decades of research have identified hundreds of genes who disruption is associated with the development of CHDs [18]. However, many studies designed to better understand the causative mechanisms underlying CHDs often overlook any contribution from disruption to the extraembryonic tissues and placenta. In the present study, we characterized fetal and placental development at term in the Ohia mouse line which displays an HLHS-like phenotype. Most notably, no double homozygote fetuses were recovered at GD18.5 and further analysis of the placentas carrying the heterozygous mutation revealed a shifted frequency towards lower fetal weight. This shift in fetal weight was associated with changes to the placenta microstructure; reduced labyrinth area and fetal capillary volume density, and reduced gene expression of angiogenic factors indicating disrupted angiogenesis within the placenta. Overall highlighting that genetic mutations which cause the disruption to fetal heart development also significantly impacts placental development.

Early in the first trimester of pregnancy, the placenta and heart develop concurrently and importantly, both organs share key developmental pathways [19-24]. During this period of development, the cellular and signaling events will ultimately determine the fate of the heart and the placenta, hence it is reasonable to assume that disrupted gene expression during first trimester development that affects one organ is likely to affect the other as genetic perturbations are infrequently restricted to one organ. Homozygous knockout of both *Sap130* and *Pcdha9* was previously shown to result in embryonic lethality before or at mid-gestation [11] but that study did not include analysis of the extra-embryonic tissues. In our study, no homozygous fetuses remained in late gestation, an observation likely explained, by placentation failure which leads to embryonic lethality [14], but beyond the current scope of this study.

Several studies in human pregnancies with fetal HLHS have shown, using ultrasound, normal umbilical artery blood flow [6, 25, 26] suggesting FGR is not due to abnormal placental blood flow and may be due to disrupted placental structure and function. A reduction in fetal capillary density in the placentas of the double heterozygotes, as well as reduced mRNA expression of *Pgf, Flt1* and *Kdr*, would, in addition, suggest abnormal placental angiogenesis. These results are supportive of outcomes in human placentas from pregnancies in which the fetus developed HLHS that show reduced mRNA expression of *PGF*, reduced protein expression of KDR, and abnormalities in both the syncytiotrophoblast and the villous vasculature [27]. *Flt*^*-/-*^ mice are embryonically lethal due to the inability of endothelial cells to form organized vascular channels in early embryonic development, including in the extraembryonic tissue [28] further highlighting the role for this gene in vascular development. Overall, such findings would indicate an imbalance in the angiogenic mechanisms within the placenta that may contribute to the increased frequency of lower weight fetuses carrying the heterozygous genotype.

One of the predominant roles of the placenta is to coordinate the transport of nutrients from mother to fetus [9]. We observed positive correlation between nutrient transporter mRNA expression and fetal weight in the double heterozygotes, which was lacking in the wildtype genotype, suggesting a possible compensatory response of the placenta to the maldevelopment/malfunction. Additionally, protein expression of amino acid transporters Slc7a5 and Slc7a8 was increased in the double heterozygote placentas compared to wildtype. In the placenta, Slc7a5 and Slc7a8 coordinate the uptake and transport of large neutral amino acids crucial for fetal growth and development [29]. Analysis of human placentas from pregnancies complicated by HLHS also show disrupted protein expression of Slc7a5 and Slc7a8, although this was shown to be lower in the HLHS placentas compared to normal controls [30]. Never-the-less such findings corroborate with the hypothesis that the mechanisms that underlie the development of CHDs also impact placental nutrient transporter expression.

The aim of the current study was to assess placental development and the impact on fetal growth in the Ohia mouse model which is characterized by the development of fetal cardiac defects similar to HLHS in humans. No fetuses with a double homozygous phenotype were found at GD18.5; possibly due to early placental failure that requires further investigation. This hypothesis is supported by a shift in fetal weight distribution towards lower fetal weight in fetuses that carried the double heterozygous genotype indicating perturbations in placenta function associated with the genetic mutations. The shifted fetal weight distribution was associated with reduced density of fetal capillaries in the placenta and reduced placental mRNA expression of *Pgf, Flt1* and *Kdr* which reflect similar outcomes in human cases of HLHS. Overall, whilst the double heterozygous fetuses do not carry heart defects, placental development and function is impaired; an outcome likely to be worse with the double homozygous genotype which do demonstrate a heart defect in mid-pregnancy. As such, our study highlights the usefulness of this model to continuing our understanding of the concurrent disruption of the placenta in CHDs such as HLHS.

## Declarations

### Ethic Approval

All animal procedures were performed under protocol 2015-0087 and approved by the Institutional Animal Care and Use Committee of Cincinnati Children’s Hospital Medical Center.

### Consent for publication

All authors consent for publication

### Availability of data and materials

Not applicable

### Competing Interests

The authors declare no conflicts of interest

### Funding

This work was funded by the Cincinnati Children’s Hospital and Medical Center Fetal Research Gift Fund

### Authors contributions

RLW analyzed the data and drafted the manuscript. WT, JC and AW performed animal husbandry, experiments and edited the manuscript. HNJ conceived and designed the study and edited the manuscript. All authors approve the final version.

## Notes

### Competing Interest Statement

The authors have declared no competing interest.

### Summary of Updates

Addition of new data regarding protein expression of nutrient transporters in the placenta

