## Supplemental Figure 1 for "Placental and fetal characteristics of the Ohia mouse line recapitulate outcomes in human hypoplastic left heart syndrome"

**Supplemental Material**

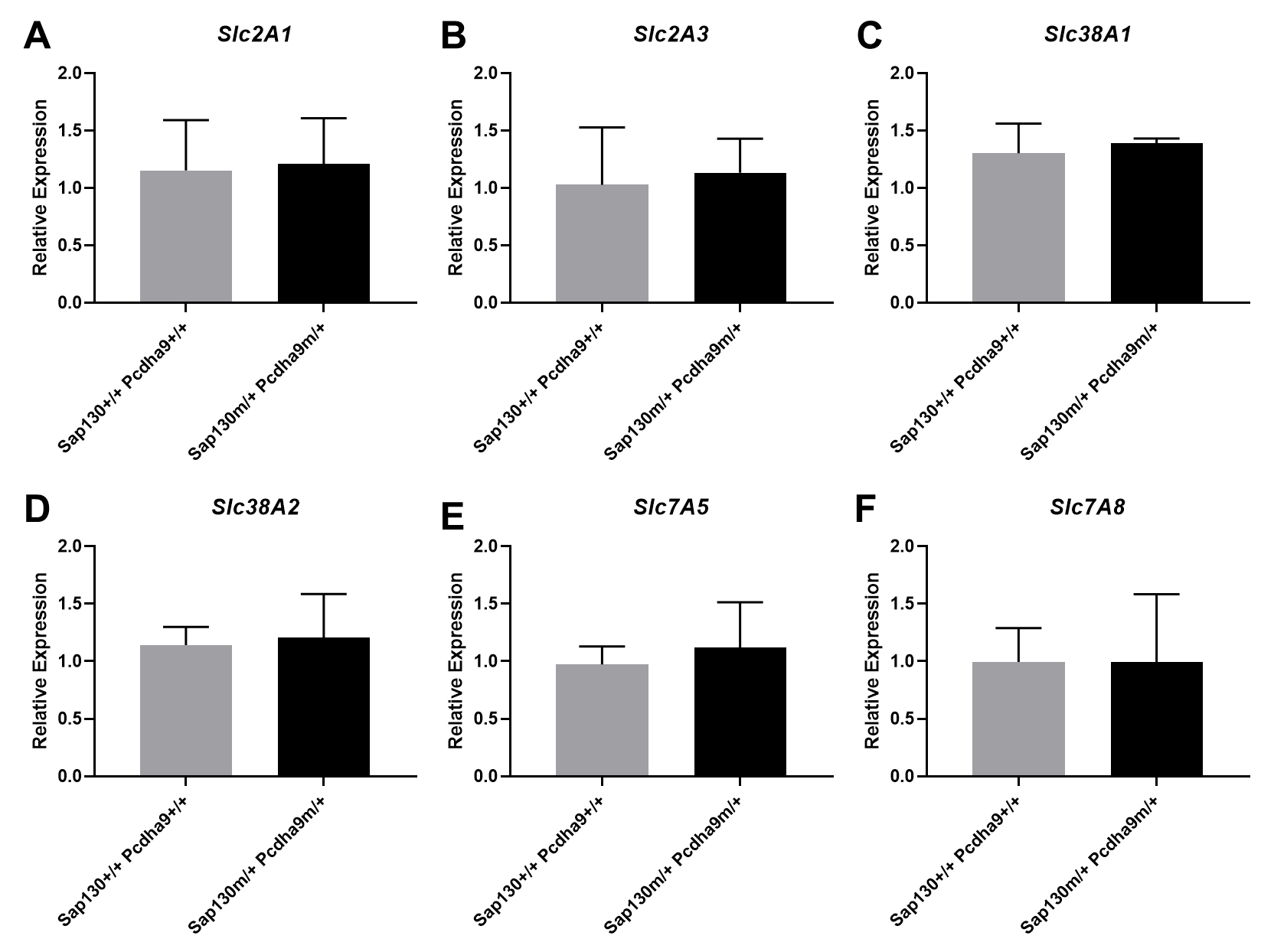

**Supplemental Figure 1** mRNA expression of nutrient transporters in the placenta of the Ohia mouse. There was no difference in expression of *Slc2A1* (**A**), *Slc2A3* (**B**), *Slc38A1* (**C**), *Slc38A2* (**D**), *Slc7A5* (**E**) and *Slc7A8* (**F**) between wildtype (*Sap130^+/+^ Pcdha9^+/+^*) and double heterozygous (*Sap130^-/+^ Pcdha9^-/+^*) placenta. Data are estimated marginal mean + standard error, n=3 females (6 wildtype and 7 double heterozygous placentas), analyzed using generalized linear modelling and including litter size as a covariate.
